# Targeted mutagenesis of Medicago truncatula Nodule-specific Cysteine-rich (NCR) genes using the Agrobacterium rhizogenes-mediated CRISPR/Cas9 system

**DOI:** 10.1101/2023.08.23.554415

**Authors:** Berivan Güngör, János Barnabás Biró, Ágota Domonkos, Beatrix Horváth, Péter Kaló

## Abstract

The host-produced nodule specific cysteine-rich (NCR) peptides control the terminal differentiation of endosymbiotic rhizobia in the nodules of IRLC legumes. Although the Medicago truncatula genome encodes about 700 NCR peptides, only few of them have been proved to be crucial for nitrogen-fixing symbiosis. In this study, we applied the CRISPR/Cas9 gene editing technology to generate knock-out mutants of NCR genes for which no genetic or functional data were previously available. We have developed a workflow to analyse the mutation and the symbiotic phenotype of individual nodules formed on Agrobacterium rhizogenes-mediated transgenic hairy roots. The selected NCR genes were successfully edited by the CRISPR/Cas9 system and nodules formed on knockout hairy roots showed wild type phenotype indicating that peptides NCR068, NCR089, NCR128 and NCR161 are not essential for symbiosis between M. truncatula Jemalong and Sinorhizobium medicae WSM419. We regenerated stable mutants edited for the NCR068 from hairy roots obtained from A. rhizogenes-mediated transformation. The analysis of the symbiotic phenotype of stable ncr068 mutants showed that peptide NCR068 is not required for symbiosis with S. meliloti strains 2011 and FSM-MA either. Our study reports that gene editing can help to elicit the role of particular NCRs in symbiotic nitrogen fixation.

## Introduction

Most legumes establish nitrogen-fixing symbiotic interaction with soil bacteria termed rhizobia. Compatible interaction between the host legumes and their rhizobia partner induces the formation of a specialised organ on the root, referred to as nodules. Rhizobia colonize the symbiotic cells of the nodules and differentiate into their symbiotic form, the bacteroids which convert the atmospheric nitrogen into ammonia^1^. The indeterminate nodules formed on the roots of temperate legumes, such as the model legume *Medicago truncatula*, possess a persistent apical meristem^2^. The continuous activity of the meristem (zone I) produces cells creating a developmental gradient in the nodules, whereupon these nodules are cylindrical in shape. The new cells mature, many of them become infected in zone II (infection zone) wherein bacteria are engulfed by a plant-derived membrane forming new organelles, the symbiosomes. Infected host cells and rhizobia differentiate in interzone (IZ) and the reduction of nitrogen takes place in zone III (nitrogen fixation zone). The completely developed and effectively nitrogen-fixing nodules produce leghemoglobin which gives wild type nodules their pink colour.

Rhizobia in the nodules of legumes belonging to the inverted repeat–lacking clade (IRLC), including *M. truncatula*, and some Dalbergoid species undergo irreversible differentiation mediated by nodule-specific cysteine-rich (NCR) peptides^3–5^. These host produced peptides contain four or six cysteine residues in conserved positions and possess signal peptides which target them to the symbiosomes through a nodule-specific protein secretory pathway^6,7^ where they govern the transition of rhizobia into bacteroids. The *M. truncatula* genome encodes a large family of NCR peptides with more than 700 members^8,9^. Although these high number of peptides show remarkable amino acid sequence variation, their function was initially presumed redundant. Nevertheless, recent studies have identified some peptides, NCR169, NCR211, NCR247, NCR343 and NCR-new35, crucial for the bacteroid differentiation and persistence in *M. truncatula* nodules but also found peptides (NCR178, NCR341, NCR344 and NCR345) which are not essential for nitrogen-fixing symbiosis^10–15^. The identification of most of these essential peptides was random during the forward genetic analyses of mutagenized *M. truncatula* populations. Therefore, the reverse genetic approach provides a comprehensive and targeted way to analyse the function of the *NCR* gene family members. The recently developed CRISPR/Cas9 (Clustered Regularly Interspaced Short Palindromic Repeats/CRISPR-associated protein 9) system is a simple and precise tool for targeted gene editing and analyse gene function in various plants^16^. Several reports described the application of CRISPR/Cas9 gene editing system in legume species (soybean^17^; peanut^18^; pea^19^; *Lotus japonicus*^20^ including *M. truncatula*^21–24^. Unfortunately, most legume species are largely recalcitrant to somatic embryogenesis and *in vitro* plant regeneration, therefore generating stable transformant legume plants using the *Agrobacterium tumefaciens*-mediated transformation system is a cumbersome and time-consuming process. In contrast, the *A. rhizogenes*-mediated hairy root transformation does not need complete plant regeneration, therefore it provides an efficient tool for a fast and large-scale functional analysis of root-related traits, such as symbiotic nodule development. Despite its robustness, the *A. rhizogenes*-mediated hairy root transformation has some disadvantages. For instance, the transformation efficiency varies, and gene transfer is not achieved in all root cells^25^, or the emerged hairy roots develop from multiple cells with different gene editing events which can result in mosaicism of transformed and untransformed cells or even multiple genotypes. This could cause trouble during gene editing of a wild-type genome to generate knockouts because a small amount of residual wild-type cells could affect the phenotype to be analysed. Therefore, the removal of untransformed or partially transformed roots and analysing the genotype of the whole hairy root is particularly important.

The CRISPR/Cas9 gene editing system, have been applied successfully to knockout and analyse the requirement of peptides NCR247, NFS1 and NFS2^12,26,27^ using *A. rhizogenes*- mediated hairy root transformation. In these reports, the targeted *NCR* genes were selected based on preliminary genetic or functional studies. In this study, we optimized the CRISPR/Cas9 system using hairy root transformation to generate mutation in selected *NCR* genes and developed a workflow to conduct their functional analysis for symbiosis.

## Results

### Testing the efficiency of a modified vector for CRISPR/Cas9 gene editing system to target *NCR169*

To analyse the function of *NCR* genes, the CRISPR/Cas9 gene editing technology was used to generate composite *Medicago truncatula* plants mutated in hairy roots. In order to monitor the presence of CRISPR constructs, delivered into the genome of *M. truncatula* root cells using the *Agrobacterium rhizogenes*-mediated hairy-root system^28^, the gene coding for the DsRed fluorescent transformation marker was introduced into the CRISPR/Cas9 plant transformation vector pKSE401 resulting in the vector pKSE401-RR. The constitutively active red fluorescent marker allowed us to remove non-transgenic roots at the early stages to promote the growth of transgenic hairy roots. To test the efficiency of targeted mutagenesis of pKSE401-RR, we designed three different single guide RNA (sgRNA) constructs targeting the *NCR169* gene, previously proved to be essential for effective nitrogen-fixing symbiosis^10^. Each sgRNA (NCR169a, b and c) was designed directing to the first half of the coding sequence of *NCR169* but the initial 5’ nucleotides for the protospacer sequences of NCR169b and NCR169c were C instead of the optimal G(/A) which is preferred by the AtU6 promoter (Fig. S1a;^29^). Nevertheless, we used these two suboptimal sgRNA constructs as well because the short, 189 bp long coding sequence of *NCR169* constrained the available optimal target sequences. The three constructs carrying each sgRNAs were introduced into roots of wild-type *M. truncatula* 2HA genotype using *A. rhizogenes*-mediated hairy root transformation. At least ten transformed roots for each construct, detected by the DsRed fluorescent marker, were sampled and used for genomic DNA isolation. The region surrounding the targeted sequence in *NCR169* was amplified by PCR. The sequence analysis of the amplicons detected only wild-type genotypes on all the NCR169b-transformed roots and roots transformed with the construct NCR169c were heterogeneous of wild-type and various mutant genotypes. However, the construct NCR169a generated mutations at higher efficiency and beside chimeric roots having both wild-type and distinct mutations, we could identify roots with homozygous or biallelic mutations for *NCR169* (Fig. S1d). The results indicated that the presence of G at the 5’ end of the protospacer sequence (G(/A)N19NGG) is crucial and using C in this position greatly reduced the efficiency of genome editing which restricts the number of possible target sites in the case of small genes such as *NCR*s. In addition, the sequence analysis of amplicons containing the targeted region revealed that the sequences obtained by Sanger method composed of multiplex sequence traces around the break site (Fig. S1b). An online tool, TIDE, enabling the decomposition of mixed sequence signals was applied^30^ that predicted the types of the mutation and the size of the deletion or insertion. For instance, the decomposition of one of the amplicons of NCR169c disclosed the mixture of wild-type and CRISPR/Cas9 edited sequences containing a 7 bp deletion and/or a one bp insertion (Fig. S1c).

In agreement with a previous report^22^, we also found that different genome editing events occurred in distinct hairy roots of a single plant which necessitates the analysis of individual nodules separately. These findings led us to the conclusions that the effectiveness of genome editing and the reliability of the identification of the nature of genome editing events needed to be improved in our system. Therefore (i) we replaced the Arabidopsis U6-26 promoter with the *M. truncatula* U6 (MtU6.6) promoter driving sgRNA synthesis, (ii) we developed a preparation procedure allowing phenotypic and genotypic analysis of individual nodules, and (iii) applied an NGS amplicon sequencing approach to assess allele distribution in individual nodules.

### Inducing mutations in *NCR169* and *NCR-new35* using the optimized CRISPR/Cas9- mediated genome editing system

Previous studies found that the *M. truncatula* U6 promoter enhanced the expression efficiency of guide or hairpin RNAs in legume species^21,31^. To increase the efficacy of targeted mutagenesis by CRISPR/Cas9, the Arabidopsis U6-26 promoter was replaced with MtU6.6 in the pKSE401-RR resulting in the vector pKSE466-RR. We inserted the sgRNAs NCR169a and another one, targeting the *NCR-new35*, of which requirement for effective symbiosis has been recently demonstrated^13^, into pKSE466-RR, respectively. To validate the efficiency of the pKSE466-RR vector construct, the empty vector and pKSE466-RR-NCR169a were introduced into wild-type *M. truncatula* 2HA plants using *A. rhizogenes*-mediated hairy root transformation and plants were inoculated with *S. medicae* WSM419. Nodules developed on DsRed fluorescent transformed roots were harvested 3-4 weeks post inoculation (wpi) with rhizobia, individually fixed and sections were stained with nucleic acid-binding dye SYTO13. Following image capturing, genomic DNA was purified from nodule sections and the regions surrounding the targeted sequences were amplified by PCR.

The elongated nodules developed on empty vector-transformed roots were pink and showed the characteristic zonation of indeterminate nodules (Fig. 1a and d). The nodules on the roots of *M. truncatula* 2HA transformed with the pKSE466-RR-NCR169a were either pink and elongated, suggesting that these nodules were functional wild-type (Fig. 2g, i and k), or white undeveloped, implying that these were ineffective symbiotic nodules (Fig. 1b and 2b, d and f). To determine the nature of the mutations at the targeted region, the sequence analysis of the amplicons, generated from randomly selected elongated and undeveloped nodules, was carried out. The targeted amplicon sequences were determined using a next-generation sequencing (NGS) approach and the quantitative allele composition and the verification of genome editing mutations were analysed using the online tool CRISPResso2^32^. The analysis revealed that all the white undeveloped nodules carried few base pair biallelic insertions or deletions at a high ratio and the very few reads of wild-type alleles identified in these nodules were probably derived from sporadic wild-type cells or the trace amount of wild-type contamination (Fig. 1b, Fig. 2a-f). The morphology and bacterial occupancy of these SYTO13-stained undeveloped nodules were very similar to the nodulation phenotype of the deletion mutant *dnf7-2* showing no fluorescence in the nodule region proximal to the root (Fig. 1e). In addition, these plants, compared with empty-vector-transformed plants, showed the symptoms of nitrogen starvation (yellowish leaves and stunted growth habit) indicating the defect of symbiotic nitrogen fixation of these nodules (Fig. 1b). Contrary to these, the sequence analysis of the randomly selected elongated pink nodules showed that they either carried almost exclusively wild-type alleles (Fig. 2g and h) or they were heterozygous for example for a 4 bp deletion and the wild-type alleles (Fig. 2i and j). We also found a wild-type nodule that predominantly carried a homozygous deletion of 3 base pairs resulting in a deletion of the first single Glu residue of the mature peptide of NCR169 which mutation did not abolish the symbiotic function of NCR169 (Fig. 2k and l).

**Figure 1.**
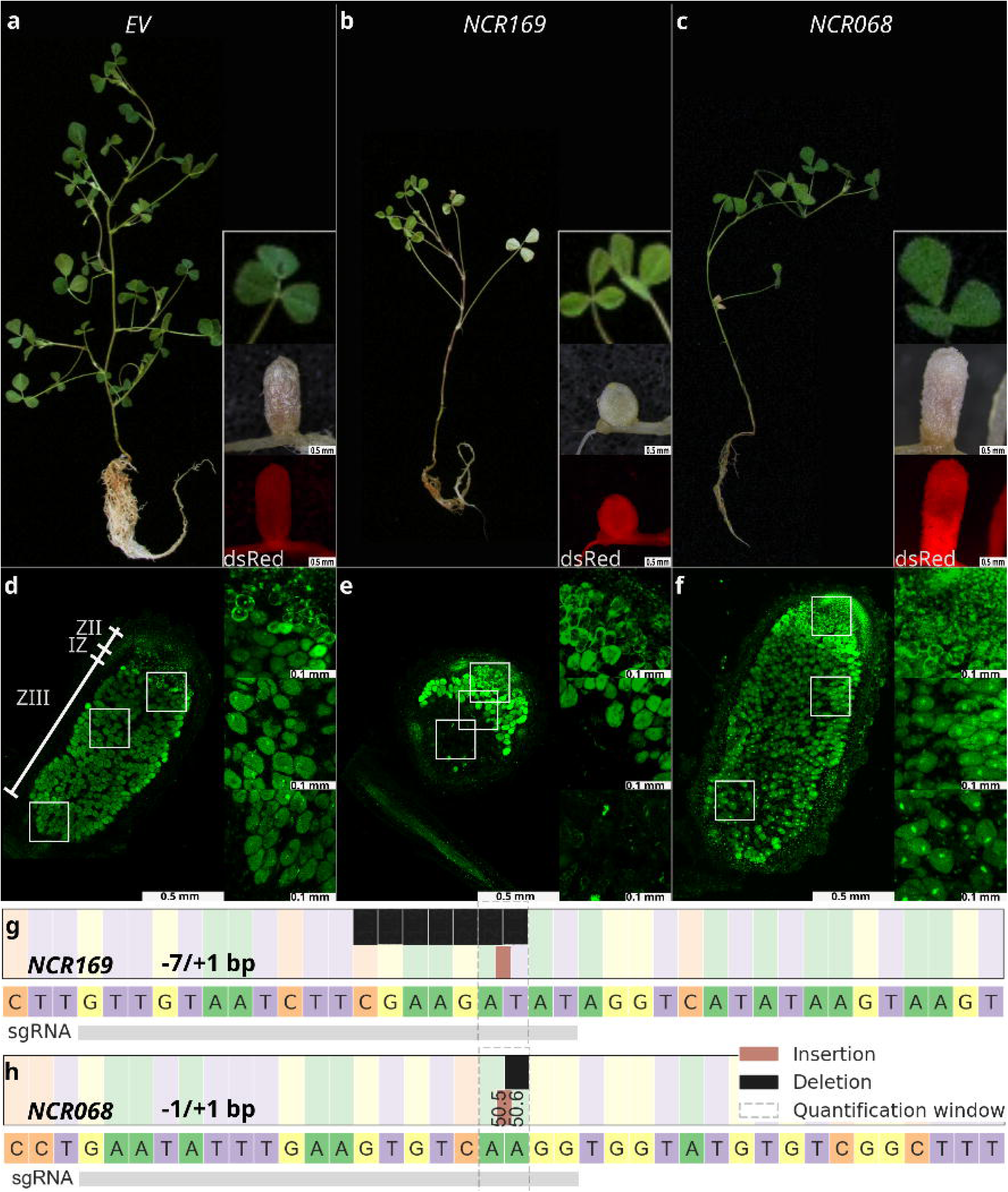
Plant gross and nodulation phenotype of *Agrobacterium rhizogenes*-mediated transgenic *M. truncatula* Jemalong 2HA hairy-roots mutagenized with CRISPR/Cas9 gene editing. No sign of nitrogen deficiency was observed on plants transformed with empty vector (**a**) or the construct targeting *NCR068* (**c**) 4 weeks post inoculation with *S. medicae* WSM419. Transgenic roots were identified by fluorescence of DsRed protein. Elongated pink nodules, showing the characteristic features of indeterminate nodules, developed on transgenic roots. These nodules were fully colonized by rhizobia visualized by the fluorescent dye SYT013 (**d** and **f**). (**b**) Targeted mutagenesis of *NCR169* resulted in transgenic plants showing the symptoms of nitrogen starvation. Small, spherical and white ineffective nodules were formed on *NCR169-*targeted roots. (**e**) *ncr169* mutant nodules did not contain infected cells in the region corresponding to the nitrogen fixation zone of wild-type plants. Inset images in (**d-f**) show the higher magnification of cells in nodule regions indicated by white rectangles. (**g** and **h**) The targeted regions were amplified and sequenced from the transgenic nodules. The sequence analysis revealed biallelic mutations causing frameshifts in edited *NCR169* (g) and *NCR068* (h) genes. ZII: infection zone; IZ: interzone; ZIII: nitrogen fixation zone; EV, empty vector

**Figure 2.**
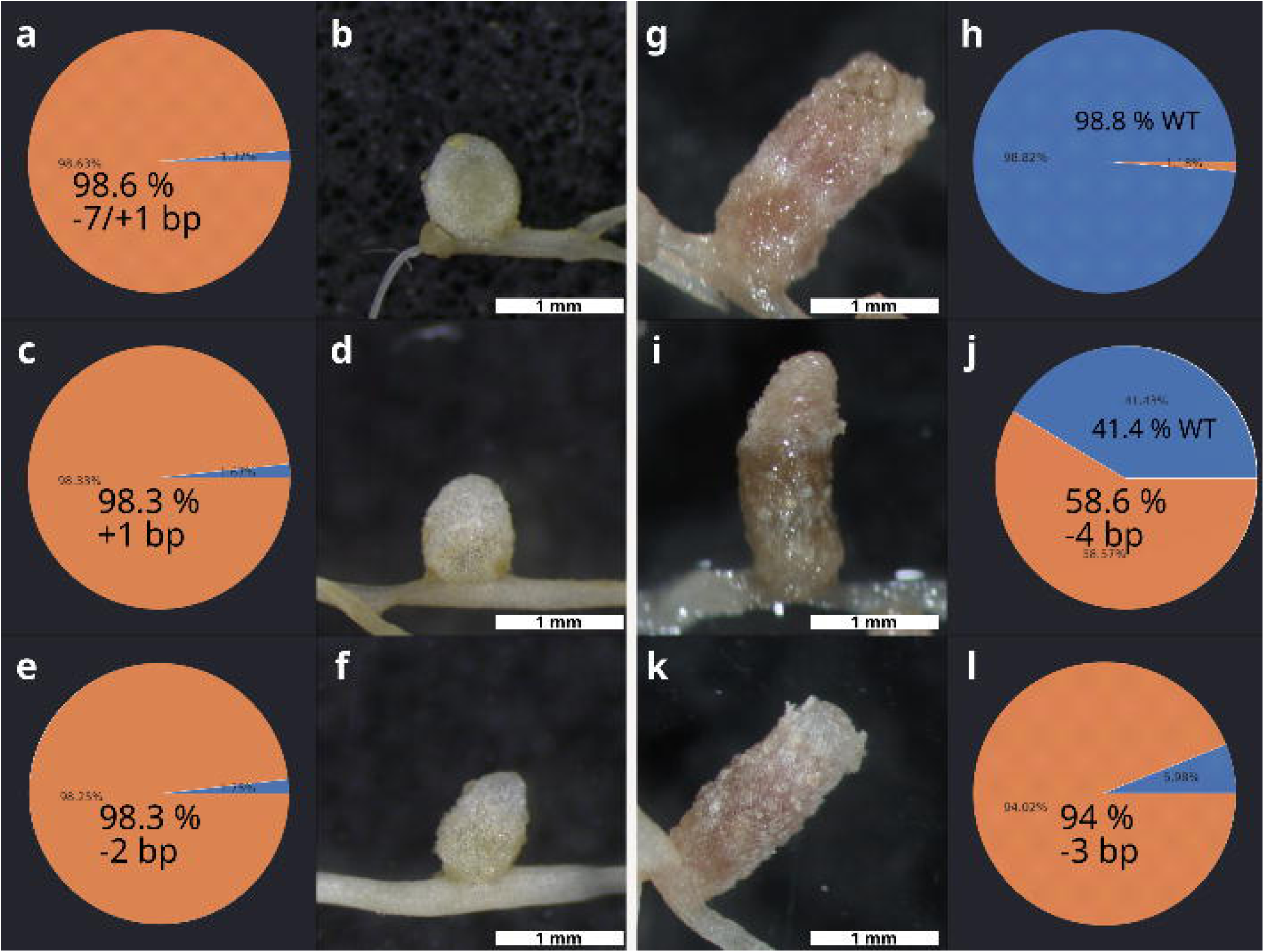
The nodulation phenotype of hairy roots transformed with the construct pKSE466- RR-MtNCR169a was associated with the proportion of wild-type *NCR169* allele or with the presence of mutant NCR169 lacking a single residue. White (Fix-) and pink (Fix+) nodules were randomly harvested 3-5 wpi with *S. medicae* WSM419 and the result of gene editing was identified by the sequencing of the targeted region using an Illumina platform. The ratios of reads assigned to mutant (brown ochre) and wild-type (blue) alleles are presented in pie chart. Fix- nodules (**b**, **d**, and **f**) carried almost exclusively insertions or deletions resulting in frameshift in the coding sequence of *NCR169* (**a**, **c** and **e**). Contrary, pink nodules (**g**, **i** and **k**) carried either unmodified alleles on a large scale (>40%) (**h** and **j**) or a deletion of 3 base pairs that did not result in the loss of function of NCR169 (**l**).

We also applied the developed CRISPR/Cas9 toolkit to target *NCR-new35*, a recently identified *NCR* gene which is crucial for the persistence of bacteroids in *M. truncatula* cv. Jemalong nodules^13^. The empty vector and the pKSE466-RR-NCRnew35 constructs were introduced into the roots of *M. truncatula* 2HA and *M. truncatula* ecotype R108 plants using *A. rhizogenes*-mediated hairy root transformation. Nodules on empty vector-transformed roots displayed the characteristic zonation of indeterminate nodules showing cells occupied by rhizobia (Figure S2e and m). In contrast, undeveloped nodules were observed on roots of *M. truncatula* 2HA and R108 plants transformed with the editing construct of *NCR-new35* (Figures S2 and Fig. S3g-r and Fig. 4e-h). The sequence analysis showed that undeveloped nodules, randomly collected from transformed roots of *M. truncatula* 2HA and R108 plants, carried biallelic or monoallelic homozygous mutations in the amplicons containing the target sequence in *MtNCR-new35* (Fig. 4q and r). SYTO13-staining revealed that these nodules did not show the typical zonation and only their apical part fluoresced indicating that the region corresponding to the prospective nitrogen fixation zone was devoid of rhizobia (Figure S2f-h and n-p). The morphology and bacterial occupancy of genome edited nodules were in agreement with the nodulation phenotype of *Mtsym19* and *Mtsym20* nodules defective in NCR-new35^13^. These results also revealed that NCR-new35 is crucial for the symbiotic interactions not only between *M. truncatula* 2HA and *S. medicae* WSM419 but between *M. truncatula* R108 and *S. meliloti* FSM-MA as well.

### Selecting of *NCR* genes to be targeted by CRISPR/Cas9 editing

The *M. truncatula* genome contains a large gene family coding for more than 700 NCR peptides but only few of them, NCR169, NCR211, NCR247, NCR343 and NCR-new35, have been proved to be crucial for symbiosis^10–15^. The sequences of the essential *NCR* genes are highly variable, and their common feature is that they contain four cysteine residues in conserved positions. The peptide NCR247 is unique among the crucial and most of the other NCR peptides because it contains five residues between the third and fourth conserved cysteines compared with the canonical four internal residues^13^. In addition, no common feature could be identified among few NCR peptides found to be non-essential for symbiosis^10,11,13^. Therefore, it was not straightforward to ascertain which additional NCR peptides are essential and should be targeted. Despite all that, we have set up a selection pipeline to identify *NCR* genes to be targeted. We have analysed several features of more than 700 predicted *NCR* genes identified in the *M. truncatula* genome^9^. The signal peptides of NCRs are crucial for delivering the peptides to the bacteroids^4,6,7^, therefore, (i) first we selected 655 *NCR* genes coding for peptides possessing a predicted signal peptide (Fig. 3a). The pool of *NCR* genes to be targeted (ii) were further enriched by selecting the encoded peptides with four cysteine residues in conserved positions. (iii) We have analysed the expression profile of the diminished list of 248 *NCR* genes in the database obtained by laser-capture microdissection of *M. truncatula* nodules^33^. We identified 11 *NCRs* with no reads, evidently seem to be pseudogenes, and 65 *NCRs* which are expressed at low level (< 10 000 reads) in nodules compared with other NCRs. The recently identified *NCR-new3*5 shows the lowest expression level among the crucial NCRs^13^ and thus, we have selected genes from the 172 *NCRs* showing similar or higher expression level as *NCR-new35* (Fig. 3c and d). (iv) These *NCR* genes were analysed to predict whether they had contained sequences which were amenable to CRISPR-based editing. The cysteine residues are essential for the function of NCR peptides *in planta* ^10,13^, therefore we have prioritised those *NCRs* which could be targeted prior to the fourth conserved cysteines to generate presumably non-functional peptides. We have further searched the *M. truncatula* genome using an online prediction tool CRISPOR^34^ for off-target sites that could be recognized by sgRNAs designed for *NCR* genes. The isoeletric points (pI) of crucial NCR peptides vary between 4.78 and 10.15 implying that anionic, neutral and cationic NCR peptides could be essential for nitrogen-fixing symbiosis in *M. truncatula*. For this reason, the gene of NCR peptides with wide range of charges were selected to be edited (Fig. 3b). (v) Finally, we have selected four *NCR* genes, *NCR068*, *NCR089*, *NCR128* and *NCR161*, to be targeted by CRISPR/Cas9 editing. The pI of the encoded mature peptides varied between 5.75 and 8.34 (Fig. 3b). *NCR128* and *NCR161* are induced in the late infection zone and showed expression in the interzone but *NCR068* and *NCR089* are preferentially expressed in the interzone and at lower level in the nitrogen fixation zone based on the LCM transcription data (Fig. 3c,^33^). The transcription activity of the four selected *NCRs* was lower than *NCR169* and *NCR211* but similar or higher than *NCR- new35* (Fig. 3d).

**Figure 3.**
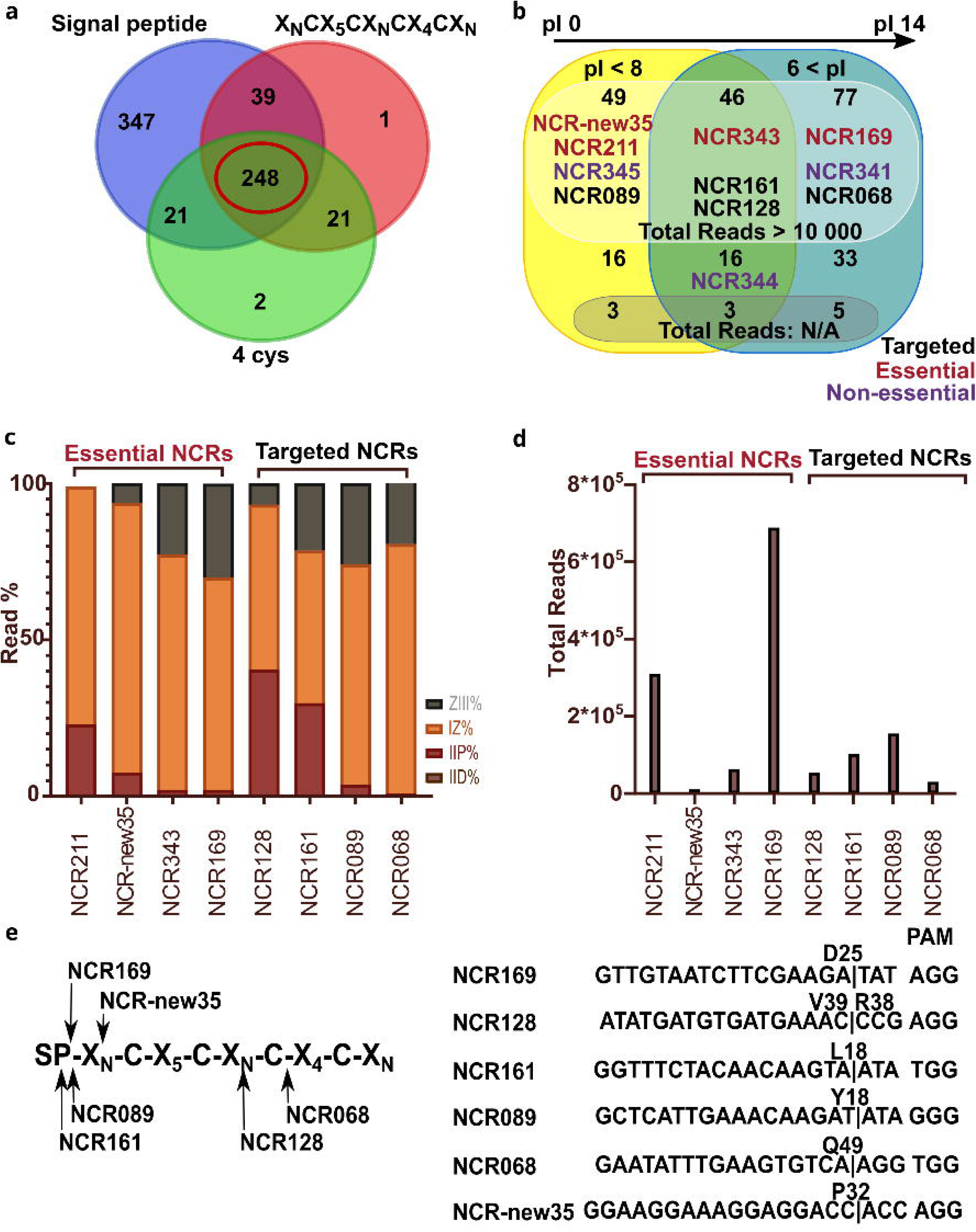
Selection of *NCR* genes to be targeted by CRISPR/Cas9 mutagenesis. (**a**) 679 *M. truncatula NCR* genes were grouped based on the presence of a signal peptide in their encoded peptide sequence and containing four cysteine residues in conserved positions. (**b**) the 248 NCR peptides complying with all the three features (**a**) were classified based on the charge of the encoded mature peptides. NCR peptides crucial for symbiosis are highlighted in red, peptides targeted in this study by gene editing are in black and peptides found to be non- essential in the former study^13^ are in purple. (**c**) Relative spatial expression of four *NCR* genes essential for symbiosis and four *NCRs* mutagenized using CRISPR/Cas9 method in this study. Expression was determined by RNAseq of nodule zones obtained with laser-capture microdissection described in the study^33^. (**d**) The total read number of the *NCR* transcripts identified by RNAseq and analysed in panel (**c**). All the four *NCR*s selected for gene editing show higher transcriptional activity than *NCR-new35* showing the lowest expression activity among the crucial *NCR* genes. (**e**) The cleavage sites of Cas9 on the sequences of *NCR* genes targeted by CRISPR/Cas6 gene editing in this study. The target sites of each guideRNA were designed upstream of the nucleotides coding for the fourth conserved cysteines of each NCR peptides. ZIII: nitrogen fixation zone; IZ: interzone; IID: distal part of infection zone; IIP: proximal part of infection zone; pI: isoelectric point

### Inducing mutations in selected *NCR* genes using the CRISPR/Cas9 system

To induce mutations in genes *NCR068*, *NCR089*, *NCR128* and *NCR161* by CRISPR/Cas9- mediated cleavage, sgRNA constructs targeting a single site of each gene were designed. The designed sgRNAs targeted either the upstream region of the gene close to the sequence coding for the first residues of the mature peptide or at least at a site prior the sequence coding for the last cysteine residue (Fig. 3e). These sgRNA constructs along with the control empty vector and the constructs targeting *NCRnew-35* and *NCR169* were introduced into *M. truncatula* 2HA roots using *A. rhizogenes*-mediated hairy root transformation. Transformed plants were inoculated with *S. medicae* WSM419 at 2-3 weeks after removal of non-transformed roots and the symbiotic phenotype of transformed plants were scored at 4-7 wpi. The aerial part of plants transformed with empty vector or the constructs targeting *NCR068* did not show the symptoms of nitrogen deficiency (Fig. 1a and c) and plants transformed with constructs targeting genes *NCR089*, *NCR128* and *NCR161* displayed the similar growth habit (data not shown) indicating the effective symbiotic nitrogen fixation capacity of these plants. Nodules formed on roots transformed with sgRNA constructs of the four selected *NCRs* were elongated and pink suggesting that they were functional nodules (Fig. 4j, l, n and p). The staining of nodule sections with SYTO13 revealed that the bacterial colonization of nodules targeted for mutagenesis of genes *NCR068, NCR089*, *NCR128* and *NCR161* was identical to the wild-type indeterminate nodules developed on empty-vector transformed roots (Fig. 4i-p). Contrary, 2HA plants carrying mutations in *NCR169* or *NCR-new35* in each hairy root displayed retarded growth, showed the symptoms of nitrogen starvation characteristically (Fig. 1b) and slightly elongated white nodules were found on their roots (Fig. 4f, Figures S2 and S3). In addition, the occupancy of nodules mutagenized for *NCR169* and *NCRnew-35* showed the phenotype characteristic for the previously described deletion mutants^10,13^ (Fig. 1b, 4e-h, Figures S2 and S3).

**Figure 4.**
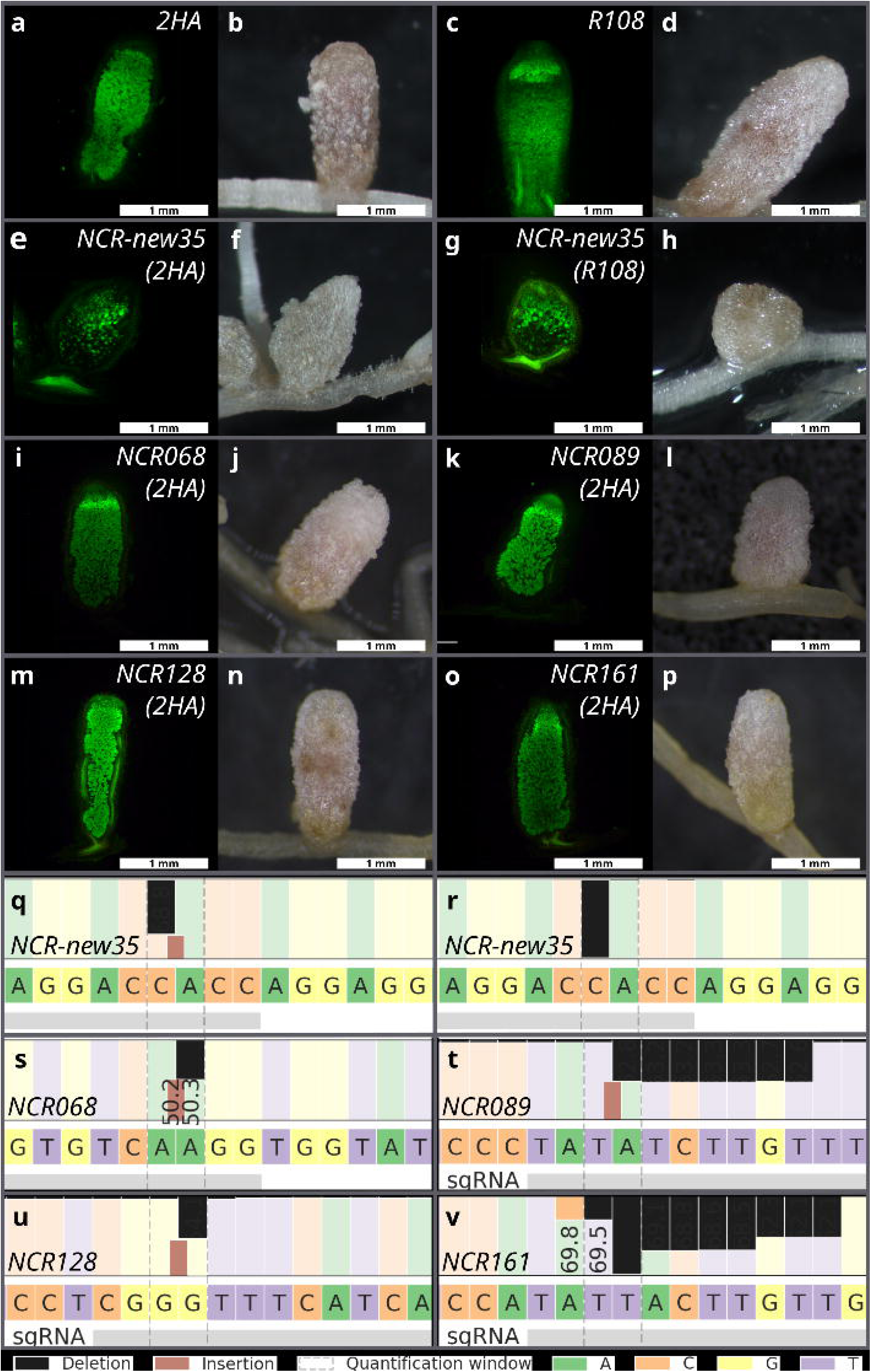
**(a-p)** Nodulation phenotype of *M. truncatula* 2HA (**e, f, i-p**) and R108 (**g** and **h**) plants mutagenized using the CRISPR/Cas9 method with *A. rhizogenes*-mediated hairy root transformation. Transformation with gene editing construct designed to target *NCR-new35* (**e**- **h**) resulted in ineffective white and slightly elongated nodules while the homozygous or biallelic mutant alleles of *NCR068* (**i** and **j**), *NC089* (**k** and **l**), *NCR128* (**m** and **n**) and *NCR161* (**o** and **p**) were similar to the elongated and pink nodules formed on empty vector-transformed roots of *M. truncatula* 2HA (a and b) and R108 (**c** and **d**) plants. Images of nodule sections following SYTO13 staining and whole nodules were taken 4-5 weeks post inoculation with *S. medicae* WSM419 used for *M. truncatula* 2HA and *S. meliloti* FSM-MA for R108 plants. (**q-v**) Allele-specific results of CRISPR/Cas9 editing of the targeted *NCR* genes identified by next generation sequencing on Illumine platform. Black and brown rectangles show deleted and inserted nucleotides, respectively.

To verify the result of CRISPR/Cas9 genome editing, the targeted regions were amplified from nodule sections following microscopic analysis and sequenced using a MiSeq platform. The sequence analysis of the amplicons was carried out with the online tool CRISPResso2. The Cas9-induced mutations were predominantly indels, with frequencies between 47 and 100% of total reads (Table 1.). The genotyping also revealed that individual nodules were often chimeric containing more than two alleles or the ratio of the two alleles deviated from the expected 1:1 ratio indicating that the tissues of these nodules on hairy roots originated from multiple founder cells with different genetic background. The identified indels generated frameshift mutations in the targeted *NCR* genes presumably producing malfunctioning peptides. Nevertheless, all the nodules carrying biallelic mutations in the targeted *NCRs* were elongated and pink, indicating the presence of leghemoglobin, and were fully colonized by rhizobia indicating that peptides NCR068, NCR089, NCR128 and NCR161 are not required for the effective symbiotic nitrogen fixation between *M. truncatula* 2HA and *S. medicae* WSM419 (Fig. 4i-p. and S4).

**Table 1.** Editing efficiency of different targeted *NCR* genes in nodules harvested from hairy roots mediated by *A. rhizogenes* transformation. Randomly selected nodules were harvested 3-5 wpi with *S. medicae* WSM419 and total DNA was isolated from fixed nodule sections. The PCR amplified target regions were sequenced on Illumina platform and the reads were analysed by using the CRISPresso2 tool. The gene *NCR169* was targeted for gene editing as a control. Three pinkish elongated nodules were selected randomly from hairy roots and also three small white nodules were picked randomly from plants which did not form any pink nodules on their roots. *One of the pinkish nodules carried mutant alleles on large proportion (94%). This mutation was a three bp deletion that resulted in an in frame deletion of the first Glu residue of the mature peptide NCR169. The lack a of this single residue probably did not abolish the function of peptide NCR169.

|  | NCR169 |  | NCR068 | NCR089 | NCR128 | NCR161 |
| --- | --- | --- | --- | --- | --- | --- |
| *Number of sequenced nodules* | 3 white | 3 pink | 15 | 13 | 10 | 20 |
| *Number nodules carrying max. 50% WT allele* | 3 | 2 | 15 | 13 | 9 | 19 |
| *Number of fully mutant nodules (mismatch/indel)* | 3 | 0* | 14 | 13 | 9 | 12 |
| *Number of fully mutant nodules (only indel)* | 3 | 0* | 7 | 13 | 9 | 12 |
| *Nodule samples carrying only indel alleles* | 100% | 0% | 47% | 100% | 90% | 60% |

### The nodulation phenotype of stable mutant of *NCR068* regenerated from gene edited hairy roots is not strain dependent

To analyse the symbiotic phenotype of genome editing mutants in more detail, we regenerated transgenic plants from hairy roots mutagenized for *NCR* genes. As a proof of concept, we cut root segments that had been transformed with the editing construct of *NCR068* and stable transformants were regenerated using a slightly modified formerly published protocol^35^. DNA was purified from the leaves of hairy root-derived regenerated plants and the sequence analysis revealed that all the plants were homozygous containing a single adenine insertion in *NCR068* (Fig. 5g). The one bp insertion generates a frame shift in the coding sequence of *NCR068* and the mutant sequence encodes a chimera peptide containing 27 NCR068-specific and 23 non-NCR068-specific residues before termination. The fourth cysteine residue in the conserved position is lacking in the chimeric NCR068 peptide and based on the necessity of cysteines for the function of NCRs *in planta* ^10,13^, we reckoned that the mutant NCR068 peptide does not function correctly.

**Figure 5.**
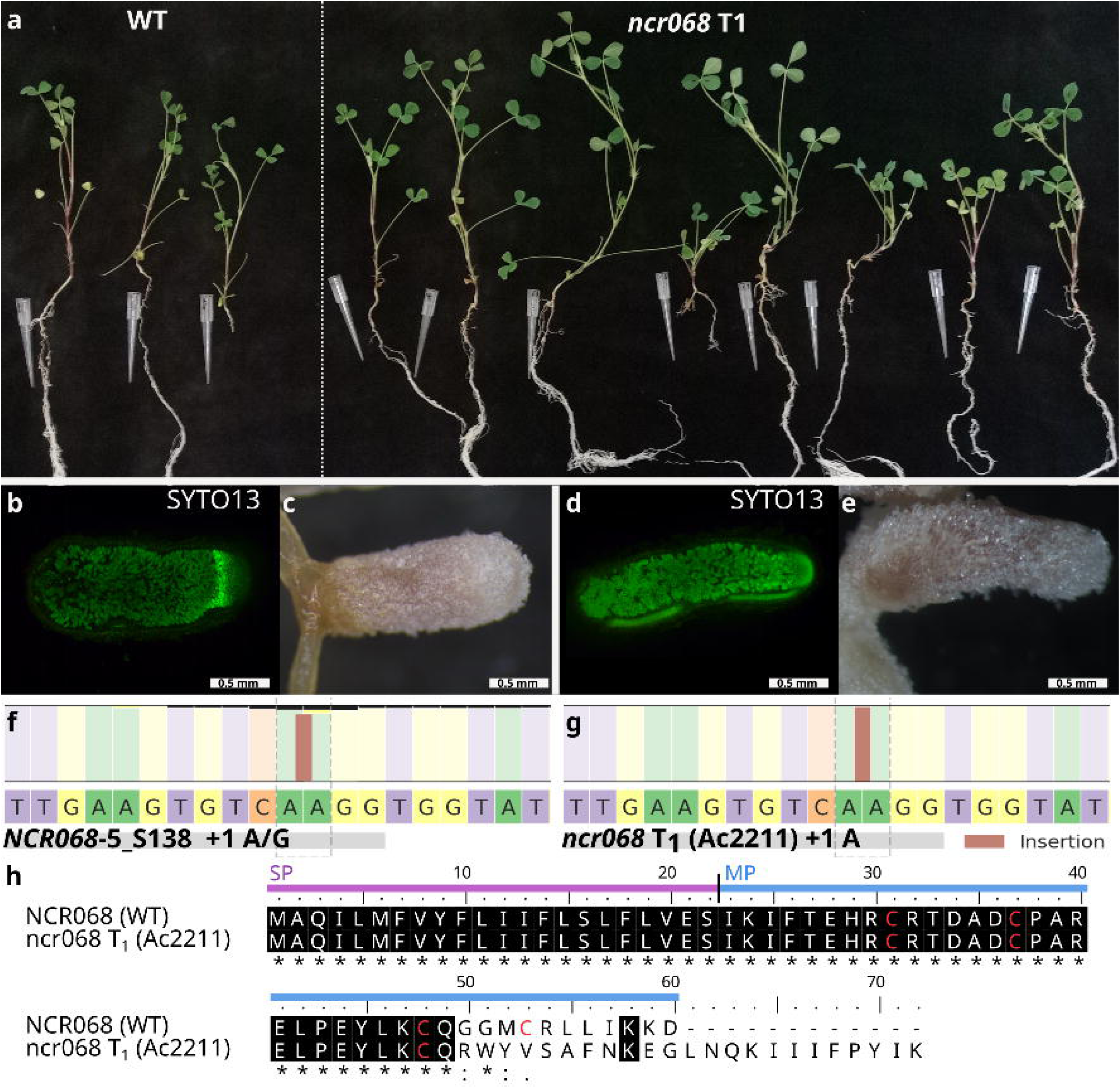
(**a**) Stable *ncr068* mutants regenerated from segments of transgenic hairy roots with induced mutations in *NCR068* using the CRISPR method retained wild type symbiotic phenotype similar wild wild-type (WT) *M. truncatula* 2HA plants 3 wpi with *S. medicae* WSM419. (**b** and **c**) The gene edited nodules developed on *A. rhizogenes*-transformed hairy-roots were elongated and pink with colonized cells in the nitrogen fixation zone. (**d** and **e**) Nodules developed on the roots of regenerated stable *ncr068* mutants were similar to the nodules found on hairy-roots mediated by *A. rhizogenes*. (**f**) The selected hairy root (NCR068-5_S138) was chimeric and composed of cells with homozygous insertions of A or G in the targeted region of *NCR068*. (**g**) The regenerated stable *ncr068* mutants (Ac2211) carried a homozygous A insertion causing a frameshift in the coding sequence of *NCR068*. (**h**) The alignment of WT and mutant NCR068 peptide sequences encoded by the mutated gene in the stable *ncr068* mutant. Identical residues are highlighted by black background, conserved Cys residues are in red. SP, signal peptide; MP, mature peptide

To confirm the symbiotic phenotype of regenerated *ncr068* mutant, T1 seeds were synchronously germinated and planted into substrate with no nitrogen content. Three wpi with *S. medicae* WSM419, *ncr068* stable transformants did not show symptoms of nitrogen starvation and developed like wild type *M. truncatula* 2HA plants (Fig. 5a). Nodules, formed on the roots of stable transformant *ncr068* mutant plants were elongated similarly what we found on the hairy roots of composite plants induced by the transformation construct targeting *NCR068* (Fig. 5c and e). The staining of *ncr068* nodule sections with SYTO13 revealed infected cells in interzone and nitrogen fixation zone fully occupied with elongated bacteria, similarly as it was found in wild-type nodules and *NCR068-*targeted hairy roots (Fig. 1d and f, 5b and d). The nodulation phenotype of the regenerated mutant *ncr068* confirmed that the peptide NCR068 is not essential for the nitrogen-fixing symbiosis between *M. truncatula* and *S. medicae* WSM419.

Different symbiotic efficiency has been reported for *Sinorhizobium* sp.^36,37^, therefore we assayed the strain-dependent nodulation phenotype of regenerated *ncr068* mutant in response to *S. meliloti* strains 2011 and FSM-MA compared with *S. medicae* WSM419. The growth habit of wild-type *M. truncatula* 2HA and *ncr068* mutant plants was similar following inoculation with any of the tested strains (Fig. 6). We observed a similar shoot dry weight of wild-type and *ncr068* plants inoculated with *S. meliloti* FSM-MA or *S. medicae* WSM419 (Figure S5). In agreement with published results^38^, the performance of wild-type and, in our study *ncr068* plants as well, inoculated with *S. medicae* WSM419 superseded the plants inoculated with *S. meliloti* 2011 (Figure S5). The *dnf7-2* mutant showed the previously described ineffective symbiotic phenotype with all three tested rhizobia strains (Fig. 6). In line with the growth habit, the nodules developed on wt and *ncr068* roots inoculated with the highly compatible strains *S. medicae* WSM419 and *S. meliloti* FSM-MA were cylindrical, pink and fully colonized by rhizobia but nodules elicited by the less effective strain *S. meliloti* 2011 were only slightly elongated (Fig. 6). In brief, no difference in plant development and nodule structure was noticed between wt and *ncr068* plants with the tested rhizobia which suggests that the symbiotic phenotype of plants lacking NCR068 is not strain-dependent.

**Figure 6.**
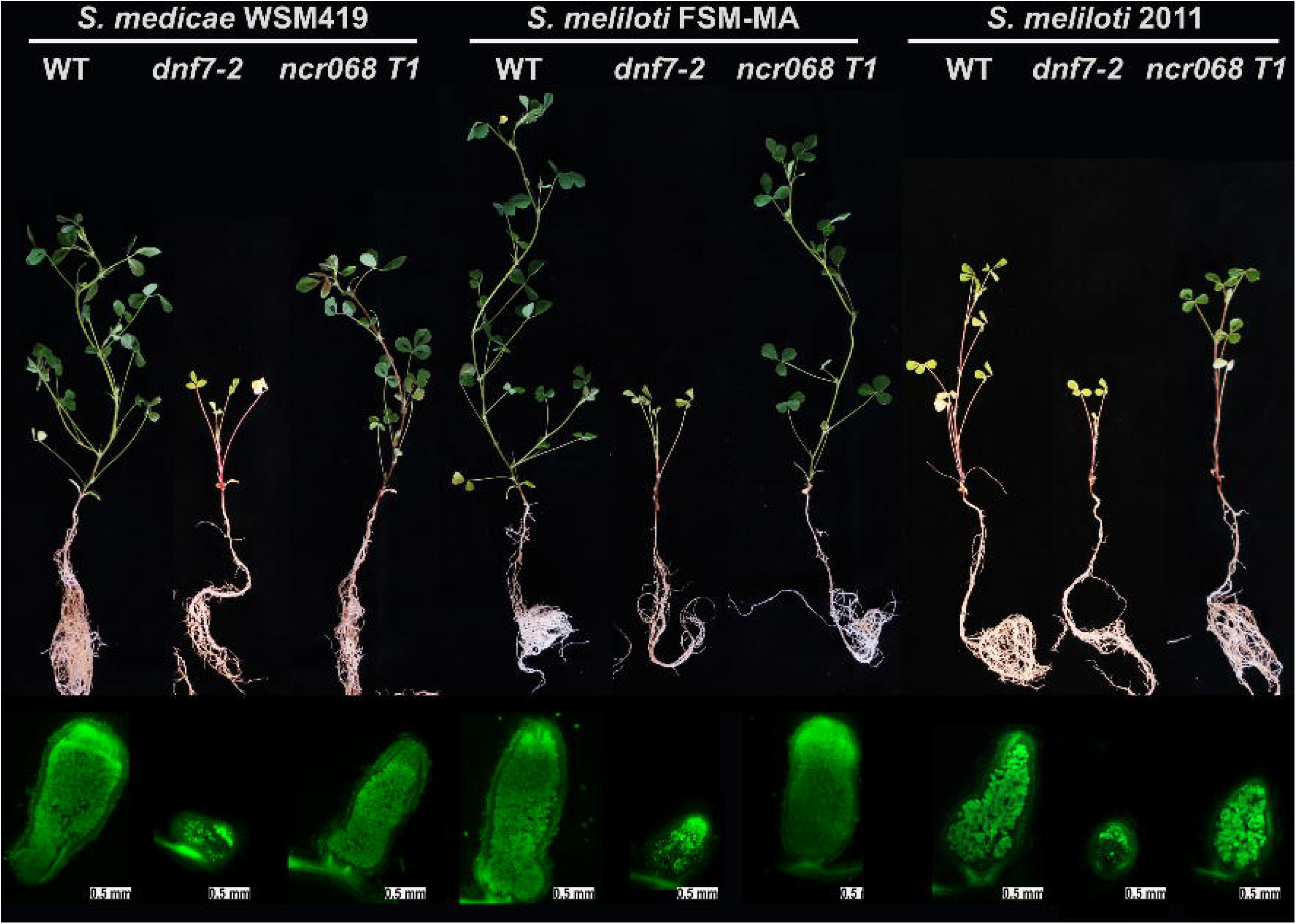
Stable *ncr068* T1 mutant plants inoculated with three rhizobia strains show wild-type symbiotic phenotype. Wild-type *M. truncatula* 2HA (WT), the deletion mutant deficient in *NCR169* (*dnf7-2*) and *ncr068* mutant plants were inoculated with *S. medicae* WSM419, *S. meliloti* FSM-MA or *S. meliloti* 2011 and scored for symbiotic phenotype at 5 wpi. WT and *ncr068* plants showed similar growth habit but presented less effective symbiotic interaction with *S. meliloti* 2011 compared with the two other strains. Mutant *dnf7-2* showed nitrogen deficiency with all three tested rhizobia strains. SYTO13-staining revealed a similar colonization of WT and *ncr068* nodules while only bacteria in infection threads and sporadic saprophytic bacteria fluoresced in the basal part of *dnf7-2* nodules.

## Discussion

NCR peptides, present solely in IRLC clade and Dalbergoid legumes, modulate the differentiation of nitrogen-fixing endosymbiont bacteria ^4,5,8^. *NCR*s compose a large gene family with more than 700 members^9^ in *M. truncatula* and because this large number of NCRs, their function was thought to be redundant. The identification of few individual NCRs crucial for nitrogen-fixing symbiosis disproved this concept ^10–15^. These essential NCRs except for NCR247, which was analysed by reverse genetic approach, were identified fortuitously using deletion mutant collections of *M. truncatula*. In addition, the large deletions in mutants *dnf4-1* and NF-FN9363 unveiled NCR peptides, NCR178, NCR341, NCR344 and NCR345, which are not crucial for the effective nitrogen-fixing symbiosis with the studied rhizobia strains^11,13^. Moreover, no specific chemical properties, charge or amino acid composition have been observed for the NCRs which would discriminate between crucial and non-crucial NCR peptides. Therefore, the targeted gene editing is the most promising tool to ascertain the need of a particular NCR peptide for nitrogen-fixing symbiosis. In this study, we used the CRISPR/Cas9 technology in the model legume *M. truncatula* to knock-out four selected *NCR* genes encoding for peptides with four conserved cysteine residues representing the anionic, neutral and cationic members of the gene family.

The *A. rhizogenes*-mediated hairy root transformation produces chimeric root system composed of different ratio of transformed and non-transformed roots depending on the efficiency of transformation. To identify transformed roots in a convenient and non-destructive way and improve gene editing efficiency, a modified vector system carrying the DsRed transformation visible marker and single-site gRNAs under the control of the *M. truncatula* U6 promoter were applied to generate mutations in the selected *NCR* genes in *M. truncatula* 2HA and R108 plants. Despite of its robustness, the *A. rhizogenes*-mediated hairy root transformation system has the disadvantage of occasionally emerged chimeric roots of a mixture of transformed and -non-transformed cells. In addition, the CRISPR/Cas9 system is not necessarily competent to edit the target sites in every cell or independent gene editing events occur in several transformed cells. Our results provided evidence of both cases; we found chimeric roots containing the mixture of wild-type and mutant alleles with different ratio, these roots mostly retained wild-type nodulation phenotype, and we detected roots having multi-allelic mutations at target sites. To eliminate these interfering effects and reveal the association between CRISPR/Cas9-induced mutations and the symbiotic phenotype without a doubt, we have sequenced the amplified fragment population of the targeted sites of SYTO13-stained and examined sections of individual nodules. This enabled the precise analysis of a large set of nodules and gene editing events could be coupled with the phenotype of each nodule.

We used two essential NCRs, NCR169 and NCR-new35, to prove the efficiency of the CRISPR/Cas9 gene editing tool in our *A. rhizogenes*-mediated hairy root transformation system and selected four NCRs (NCR068, NCR089, NCR128 and NCR161) to assay their requirement for symbiotic nitrogen fixation. We generated mutant versions of the four previously non-investigated *NCR*s and the analysis of the colonization of the corresponding nodules indicated that these four NCRs are not essential for the symbiotic interaction between *M. truncatula* 2HA and *S. medicae* WSM419. The removal of variable number of non-transgenic roots from each plant and the potential presence of root cells with non-edited genome did not enable the reliable analysis of the nitrogen starvation phenotype (stunted growth, yellow leaves) of the shoots of transformed plants. The nodules of all the previously identified mutants defective in essential *NCRs* were white and slightly elongated and showed defects in colonization of the region corresponding to the nitrogen fixation zone^10,11,13–15^. Therefore, we assessed the symbiotic phenotype of genome edited nodules mostly based on their size, colour, and the colonization of the nitrogen fixation zone. The coordinated sequence and phenotype analyses which we carried out on every single nodule, clearly demonstrated that CRISPR/Cas9-induced mutations generating corrupted open reading frames in the coding sequence of *NCR068*, *NCR089*, *NCR128* and *NCR161* did not affect the nodulation phenotype of gene edited roots, thus these NCR peptides are not essential for the nitrogen-fixing symbiosis between *M. truncatula* 2HA and *S. medicae* WSM419. Although the four target NCR genes were selected based on the common features of the encoded essential peptides (four cysteine residues in conserved positions, predicted signal peptide, nodule-specific expression, unique sequence, various pI;^10,11,13^), these criteria were not adequate to identify essentials NCR peptides. This conclusion was supported by the genetic studies that identified non-essential peptides, NCR341, NCR344 and NCR345, possessing these features. The four NCR peptides targeted in this study extend the group of NCRs proved to be non-required for nitrogen fixation under the symbiotic conditions used in this study. Our findings show that peptides NCR068, NCR089, NCR128 and NCR161 are either not crucial or have minor or redundant roles in symbiosis. Alternatively, these peptides may be required under specific symbiotic or environmental conditions or essential for strain-specific symbiotic interaction.

We have demonstrated that the *A. rhizogenes*-mediated hairy root transformation was successfully adapted to screen the requirement of NCR peptides for nitrogen fixation. We have regenerated plants of gene edited knockout of *NCR068* gene directly from transformed hairy root segments to carry out a more detailed analysis of CRISPR/Cas9-induced mutations in stable transformants. A previous proteomic study identified peptide NCR068 in bacteroids^39^ and we anticipated that NCR068 may have a crucial role in symbiosis. The mutant plants carrying a frame-shift mutation, induced by the insertion of a single base pair, in *NCR068* in their germline showed similar nodulation phenotype to nodules found on *A. rhizogenes*-mediated mutant hairy roots proving the efficiency and usability of CRISPR/Cas9- mediated mutagenesis using hairy root transformation. The stable *ncr068* mutant enabled us to assay the symbiotic phenotype with *S. medicae* strain WSM419 and two *S. meliloti* strains, 2011, a laboratory reference strain and FSM-MA, which is highly effective symbiotic partner of *M. truncatula* A17 and R108^37^. All the three tested rhizobia strains induced the formation of wild-type nodules on the root of *ncr068*, indicating that its symbiotic phenotype is not strain-specific.

Our study demonstrates that an optimized gene editing CRISPR/Cas9 vector construct and the developed workflow (Fig. S6) to analyse nodules developed on hairy roots mediated by *A. rhizogenes*- transformation is an effective system to analyse the symbiotic function of *NCR* genes. Although the selected NCR peptides were found to be non-essential for nitrogen-fixing symbiosis, the application of the technology can identify additional indispensable and crucial NCR peptides for nitrogen-fixing symbiosis.

## Materials and Methods

### Plant material, growth conditions, bacterial strains and nodulation assay

*Medicago truncatula* cv. Jemalong 2HA line^40^ (2HA) and the *M. truncatula* R108 accession (R108) were used for transformation experiments. For sterilization and scarification, seeds were soaked in concentrated sulfuric acid for 8 minutes, then rinsed with distilled water. Sterilized seeds were vernalized for 3 days on inverted 0.8 % (w/v) water agar plates at 4 °C and germinated overnight at room temperature before transformation. Seedlings were grown in zeolite substrate (Geoproduct Kft., Mád, Hungary) for five days before inoculation with rhizobia. Plants were grown in SANYO HM350 environmental test chambers at 23 °C at 16/8 hours light/dark periods.

*M. truncatula* 2HA plants were inoculated with *Sinorhizobium (Ensiferi) medicae* strain WSM419 (SmWSM419) carrying pXLGD4 plasmid^41^, constitutively expressing the *lacZ* reporter gene under the control of the *hemA* promoter. To test strain specific symbiotic phenotype of *ncr068* mutants, T1 plants were inoculated with *S. meliloti* 2011, *S. meliloti* FSM-MA and SmWSM419 (pXLGD4) strains. *M. truncatula* R108 plants were inoculated with *S. meliloti* FSM-MA. The rhizobium strains were grown overnight in liquid TA medium. The bacteria were pelleted, resuspended and diluted to OD_600_ 0.1 in nitrogen free liquid Fahraeus medium ^42^ and used for inoculation.

### Construction of the CRISPR/Cas9 binary vector used for gene editing

For CRISPR/Cas9-based gene editing experiments, a modified version of the pKSE401 vector^43^ was generated. DsRed fluorescent protein under the control of the 830 bp promoter of *UBQ10* gene from *Arabidopsis thaliana* was inserted into the *EcoR*I digested vector to allow the identification of transformed roots based on their fluorescence. This generated construct (pKSE401_RR) was further modified by replacing the AtU6 promoter with MtU6.6 promoter^31,44^ resulting the construct pKSE466_RR. The MtU6.6 fragment was amplified from the genomic DNA of *M. truncatula* A17 using the primers MtU6.6_prom_InF_F1 and MtU6.6_prom_InF_R along with the guide cassette fragment amplified from pKSE401_RR using the primers pKSE401_empty_InF_F2 and pKSE401_InF_R2 (Table S1) and the amplified fragments were assembled into the *Hind*III digested pKSE401_RR vector backbone using In-Fusion cloning (Takara Bio, Kusatsu, Japan).

### Selection of targeted genes and generating CRISPR/Cas9 constructs

To select the *M. truncatula NCR* genes to be targeted, we have analysed the previously collected and described NCR peptide sequences^9^. During the selection, first the signal peptide prediction was repeated with the SignalP 5.0 tool^45^. If the prediction did not detect signal peptide, the whole peptide sequence was considered as mature peptide. Further on, NCR peptides possessing four cysteine residues in conserved position (X_n_-C-X_5_-C-X_n_-C-X_4_-C-X_n_)^8^ were selected. The transcriptional activity of this set of *NCR* genes was analysed in the RNAseq data of *M. truncatula* nodules^33^ and genes with more than 10 000 reads were retained. The final set of 157 *NCR* genes was classified based on their isoelectric point^9^ and few genes showing low similarity to other *NCR*s were selected from each category and were selected to design sgRNA constructs. All sgRNAs were designed using the online tool available at http://crispor.tefor.net/ ^34^ using the *M. truncatula* A17 (MtrunA17r5.0) genome as reference^46^. In each case, the targeted region was in the first half of the coding sequence of the NCR gene and upstream from the last conserved cysteine residue. To minimize the risk of off-target mutations all selected sgRNA had at least two mismatches comparing with potential off-target sites (Table S2). Targets with GN_19_ NGG motif were preferred for optimal expression controlled by the MtU6.6 promoter. Plasmid constructs were assembled following the protocol described by Xing at al.^43^

### *Agrobacterium rhizogenes*-mediated hairy root transformation

CRISPR constructs were introduced into the ARqua1 strain of *Agrobacterium rhizogenes* and used for hairy root transformation of *M. truncatula* 2HA and R108 seedlings as described by Boisson-Dernier et al.^28^. Each CRISPR construct was transformed into the roots of at least 35 plants in transformation and experiments were repeated at least twice. Following transformation, the seedlings were grown for one week in square petri dishes containing antibiotic-free Fahraeus medium and then seedlings were transferred to selective Fahraeus medium supplemented with 25 mg/l kanamycin. To prevent root growth into the agar, the roots were carefully positioned between two layers of Whatman filter paper. Following the plant growth for additional two weeks, non-transgenic roots, identified based on the absence of red fluorescence, were excised. After then, plants were transferred into zeolite supplemented with Fahraeus medium containing 2 mM NH_4_NO_3_. Plants were further grown in plastic boxes to preserve rhizobium-free conditions and inoculated with rhizobia after 2-3 weeks. Prior inoculation with rhizobia, newly emerged non-transgenic roots were excised again.

### Embryonic induction and plant regeneration from hairy-root cultures

Plantlets were regenerated from transformed hairy roots grown on plate under sterile conditions using a slightly modified protocol described previously^35^. Three-week-old transgenic hairy roots were identified by DsRed fluorescence using a Leica MZ10-F fluorescence stereo microscope (Leica Microsystems GmbH, Wetzlar, Germany) and cut into approximately 0,5-1 cm segments and placed on modified MTR-2 medium (Murashige and Skoog medium including vitamins supplemented with 5.0 mg/l 2,4-Dichlorophenoxyacetic acid, 0.5 mg/l 6-Benzylaminopurine (BAP), 3 % (w/v) sucrose, 50 mg/l kanamycin, 250 mg/l cefotaxime solidified with 0.8 % (w/v) agar) to induce primary calli. After 2-3 weeks of incubation in dark at room temperature, calli were transferred to the modified MTR-3 medium supplemented with BAP (Murashige and Skoog medium including vitamins supplemented with 0.5 mg/l BAP, 2 % (w/v) sucrose, 50 mg/l kanamycin, 250 mg/l cefotaxime solidified with 0.8 % (w/v) agar) to induce embryonic calli and plates were kept in plant growing chambers with a photopheriod of 16 h of light and 8 h darkness at 24 °C. The calli were transferred to fresh medium in 1-2-week intervals. Shoots developed on MTR-3 + BAP medium were put on MTR-3 medium (without selective agent) in glass tubes and transferred to fresh medium on every 2-5 weeks to promote development and induce roots. Well-grown shoots were propagated by cutting the apical 2-3 cm of the shoots and they were transferred into separate tubes. After rooting, the T0 regenerated plants were transferred into jars containing zeolite, watered with Fahraeus medium supplemented with 2 mM NH_4_NO_3_. The jars were kept in growth chambers under the same condition as described above and humidity in the jars was gradually decreased. Once new healthy leaves emerged, the plantlets were inoculated and tested for nodulation phenotype or alternatively, they were transferred to nutrient-rich soil in the greenhouse for seed collection.

### Phenotypic characterization of transformed plants and microscopy analysis

To observe intact roots and nodules and detect fluorescence of transformed roots, a Leica MZ10F fluorescent dissecting microscope equipped with ET DsRED, and GFP LP filters was used (Leica Microsystems). Transgenic nodules were harvested, promptly immersed into ice-cold 1xPBS (pH 7.4) and fixed with 4% paraformaldehyde. Fixed nodules were embedded in 5% (w/v) agarose gel prepared in 1xPBS and 100 µm tick longitudinal sections were prepared with Leica VT 1200S vibratome. Nodule sections were stained in 1xPBS solution containing 5 µM SYTO13 dye (Thermo Fisher Scientific, Waltham, MA, USA) for 30 minutes at room temperature. Images of stained nodule sections were captured using a Leica SP5 AOBS confocal laser scanning microscope (Leica Microsystems).

### DNA isolation from sections of transgenic nodules

To purify DNA from fixed nodule sections, 2-8 sections per nodule sample were collected after SYTO13 staining and microscopic analysis and transferred into nodule lysis buffer [100 mM Tris-HCl (pH 8), 0,5M EDTA (pH 8), 0.2 % SDS, 5 mM NaCl, Proteinase K 0.02 µg/µl]. The samples were homogenized using glass pestles and incubated at 56 °C overnight. Proteinase K was inactivated by incubating the samples at 95 °C for 15 minutes. Samples were spin down by microcentrifuge and cooled them to room temperature. Subsequently 2 µL RNase A (from 10 mg/ml stock) was added and incubated at room temperature for 2 min. After that, 200 µL binding buffer [2.55M Guanidium-HCl, 10 % Triton X-100, 37 % ethanol] was added and briefly vortexed, and 200 µL of 96 % ethanol was added to the mix and vortexed immediately. The mixture was centrifuge at 8000 rpm for 30 second to pellet the tissue debris. The supernatants were then transferred to QIAprep Spin Miniprep columns (Qiagen, Hilden, Germany). Following centrifugation at 8000 rpm for 1 min, the flow-through was discarded and columns were washed two times with 500 µL washing solution (10 mM Tris-HCl pH 7.5, 80% ethanol). After the second washing step flow-through was discarded and columns were centrifuged at 13.000 rpm for 3 min. DNA elution was performed by incubating the columns with 35-50 µL of de-ionized water at 50 °C for 3 minutes and eluted DNA was collected by centrifugation at 13,000 rpm for 1 min.

### Sequence analysis of targeted genes in gene-edited nodules

The analysis of the result of target site mutagenesis of transgenic roots targeted by the sgRNAs NCR169a, NCR169b and NCR169c was performed by Sanger sequencing of PCR fragments amplified from the target region using ncr169_F1 and ncr169_R17 primer pair. To analyse the edits of targeted regions of genes NCR169, NCR-new-35, NCR068, NCR128, NCR089 and NCR161 in nodule samples, the PCR amplified fragments were analysed by next-generation sequencing (NGS). The genomic regions of appr. 200 bp, including the target sites, were amplified using GoTaq™ G2 polymerase (Promega corp. Madison WI, USA) according to the manufacturer’s instructions and gene specific primer pairs containing 5’ adaptor sequences (Table S1) The samples were sequenced on Illumina platform, generating 10 000 reads per sample. NGS sequence analysis was carried out by using CRISPresso2 program^32^ using default parameters.

## Acknowledgements

This work was supported by the Hungarian National Research Fund/National Research, Development and Innovation Office grants OTKA 119652, 129547 and 132646 as well as by the Collaborative Research Programme ICGEB Research Grant HUN17-03. We thank H. Cs. Tolnainé for her skillful technical assistance and Z. Tóth for support in confocal microscopy imaging.

## Author contributions

J.B.B. and P.K conceived the project. B.G., J.B.B., D.Á. and B.H. performed the experiments and analysed the data J.B.B. and P.K. wrote the manuscript that all authors reviewed.

## Data availability

All data supporting the findings of this study are available within the paper and the corresponding supplementary materials published online.

## Additional information

### Supplementary information

Fig. S1. Analysis of three sgRNAs targeting *NCR169*.

Fig. S2. The symbiotic phenotype of nodules formed on *M. truncatula* Jemalong and R108 hairy roots edited for *NCR-new35*.

Fig. S3. The symbiotic phenotype of nodules formed on *M. truncatula* Jemalong and R108 hairy roots edited for *NCR169* and *NCR-new35*.

Fig. S4. The nodulation phenotype of hairy roots edited for *NCR089*, *NCR0128* and *NCR161*.

Fig. S5. The shoot biomass of stable *ncr068* mutant plants inoculated with three rhizobia strains.

Fig. S6. The workflow applied for simultaneous genetic and phenotypic analysis of gene edited individual nodules.

Table S1. List of primers used in this study.

Table S2. Off-target prediction using the CRISPOR tool.

