## Supplementary Figures for "Targeted mutagenesis of Medicago truncatula Nodule-specific Cysteine-rich (NCR) genes using the Agrobacterium rhizogenes-mediated CRISPR/Cas9 system"

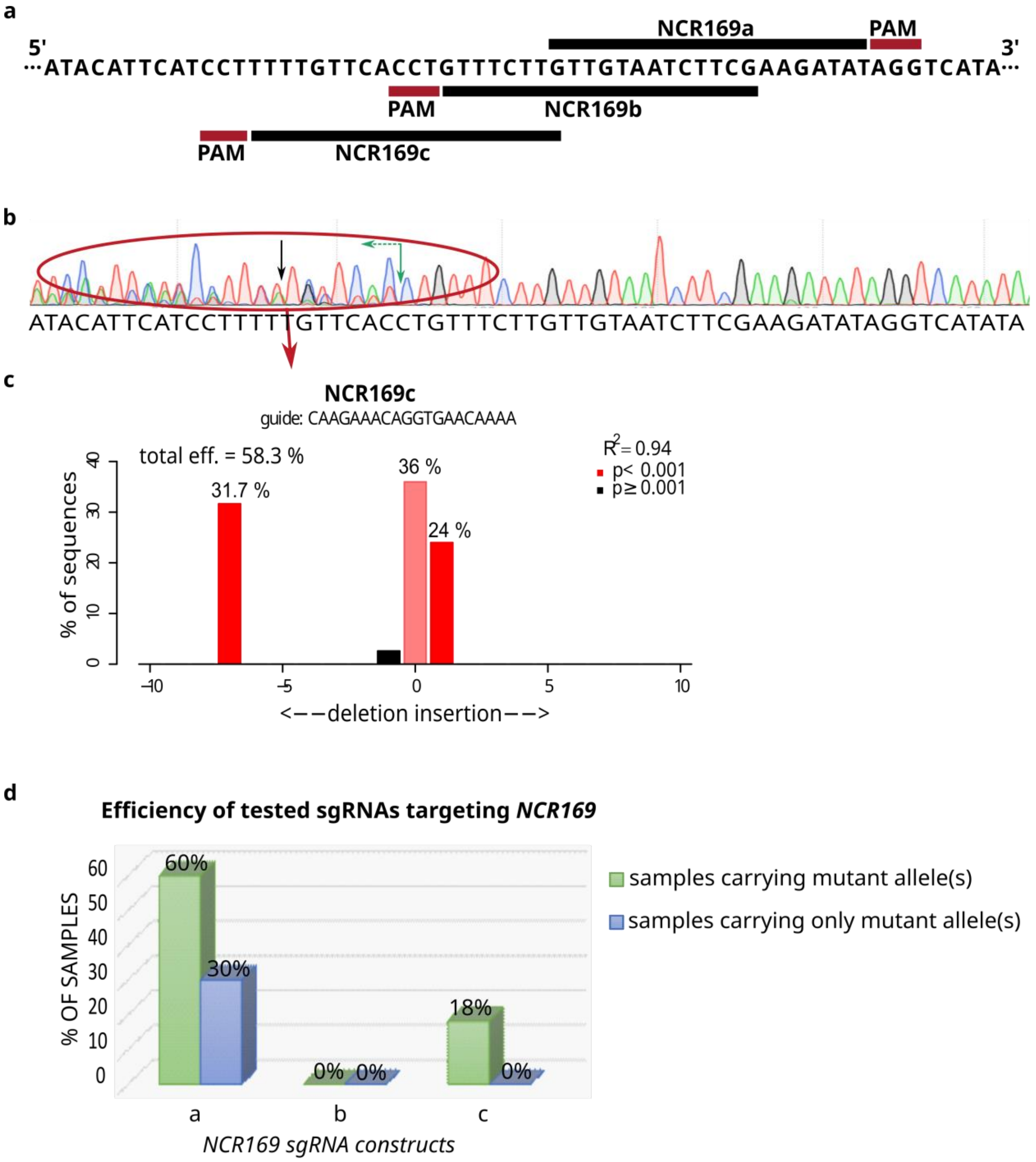


**Figure S1.** (a) sgRNAs and PAM sequences on targeted gene *NCR169*. (b) Sanger sequencing of the targeted region from a mosaic hairy root transformed with the NCR169c construct shows a mixed spectra of different edited alleles following cleavage and repair by non-homologous end joining. Black arrow shows the Cas9 nuclease-mediated double-strand break. Green arrow indicates the position of the start of mixed sequence trace. (c) The decomposition of the capillary electrophoresis sequence trace analysed by the TIDE online tool predicts wild-type alleles, a single bp insertion and a seven bp deletion in the targeted region. (d) Efficiency of the three gRNA constructs targeting the gene *NCR169*.

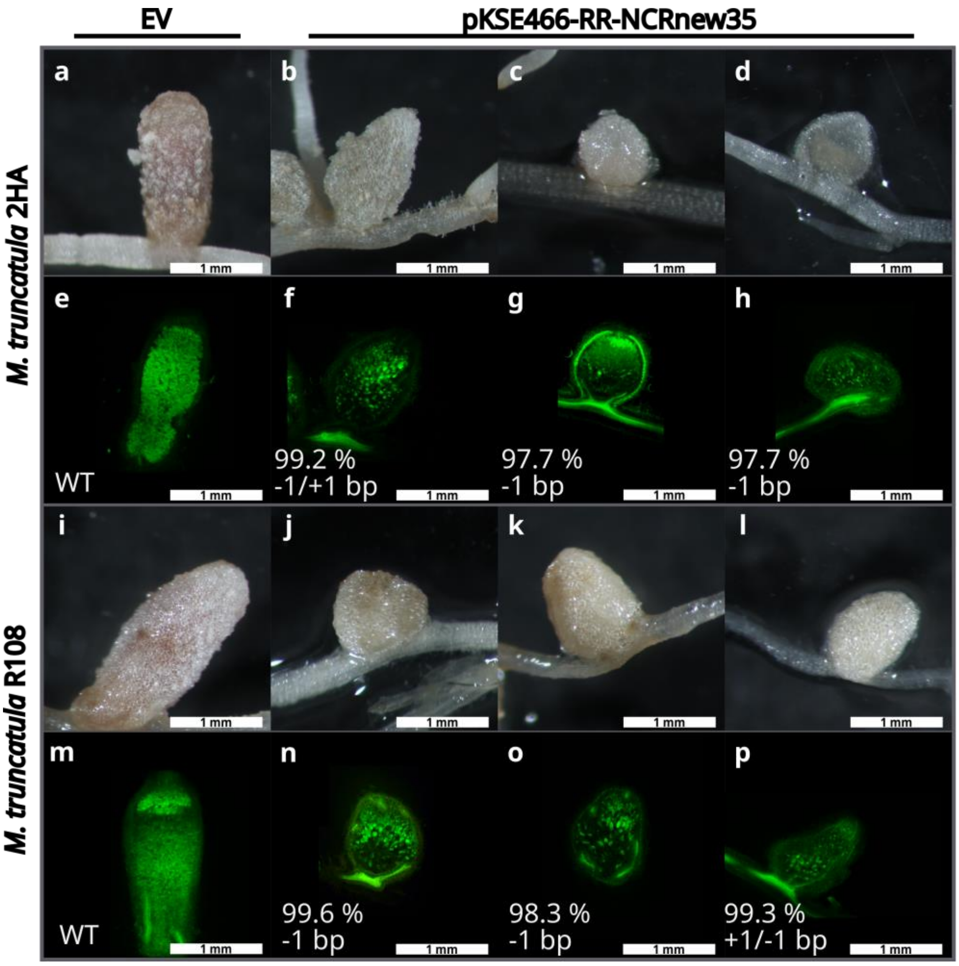


**Figure S2.** Nodulation phenotype of *M. truncatula* 2HA and *M. truncatula* R108 hairy roots mutagenized for *NCRnew35* using the CRISPR/Cas9 method with *A. rhizogenes*-mediated transformation. Nodules were fixed with paraformaldehyde, sectioned and stained with fluorescent dye SYTO13 to analyse bacterial colonization. The genotypes of the nodule sections were determined by Illumina amplicon sequencing. All nodules were harvested 3-5 weeks post inoculation with *S. medicae* WSM419 in the case of *M. truncatula* 2HA and with *S. meliloti* FSM-MA in the case of R108 plants. Nodules formed on empty vector-transformed hairy roots were pink and elongated (a and i) showing the characteristic zones of indeterminate nodules largely occupied by bacteria (e and m). The white undeveloped (b-d, j-l) and hardly colonized (f-h, n-p) nodules formed on hairy roots generated by *A. rhizogenes*-mediated transformation targeting *NCR-new35* were homozygous or biallelic for single base pair deletions and/or insertions.

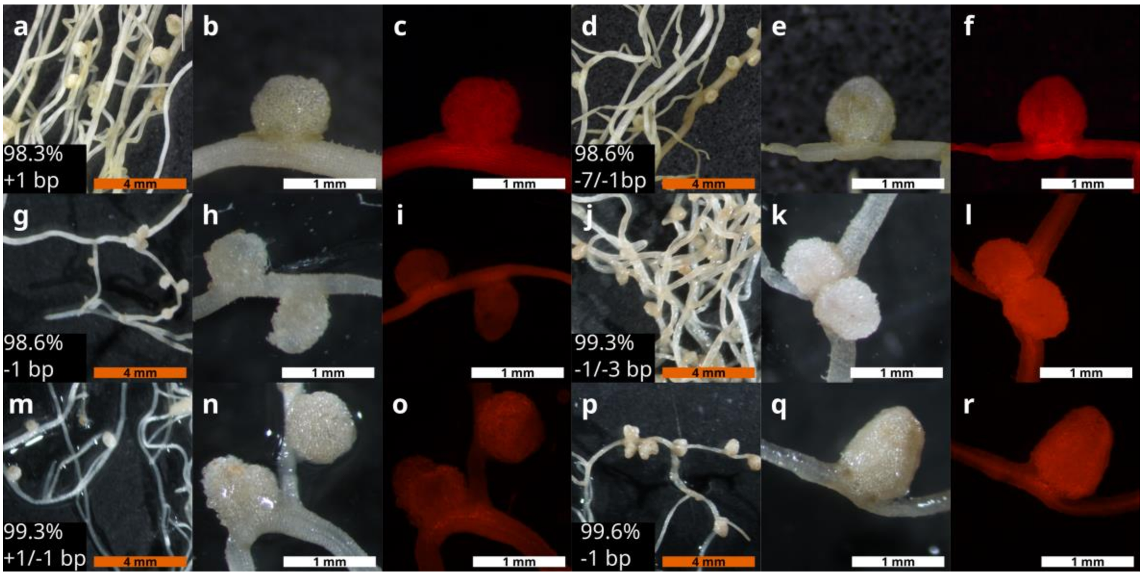


**Figure S3.** Images of hairy roots and 5-week-old nodules mutated in *NCR169* (a-f) and *NCR-new35* (g-r) of *M. truncatula* 2HA (a-l) or *M. truncatula* R108 (m-r) using the CRISPR/Cas9 method. Hairy roots generated by *A. rhizogenes*-mediated transformation were inoculated with *S. medicae* WSM419. Transgenic roots were identified by the fluorescence of the DsRed protein. The *NCR169* or *NCR-new35*-edited white undeveloped nodules carried homozygous or biallelic mutations of few base pair deletions and/or insertions.

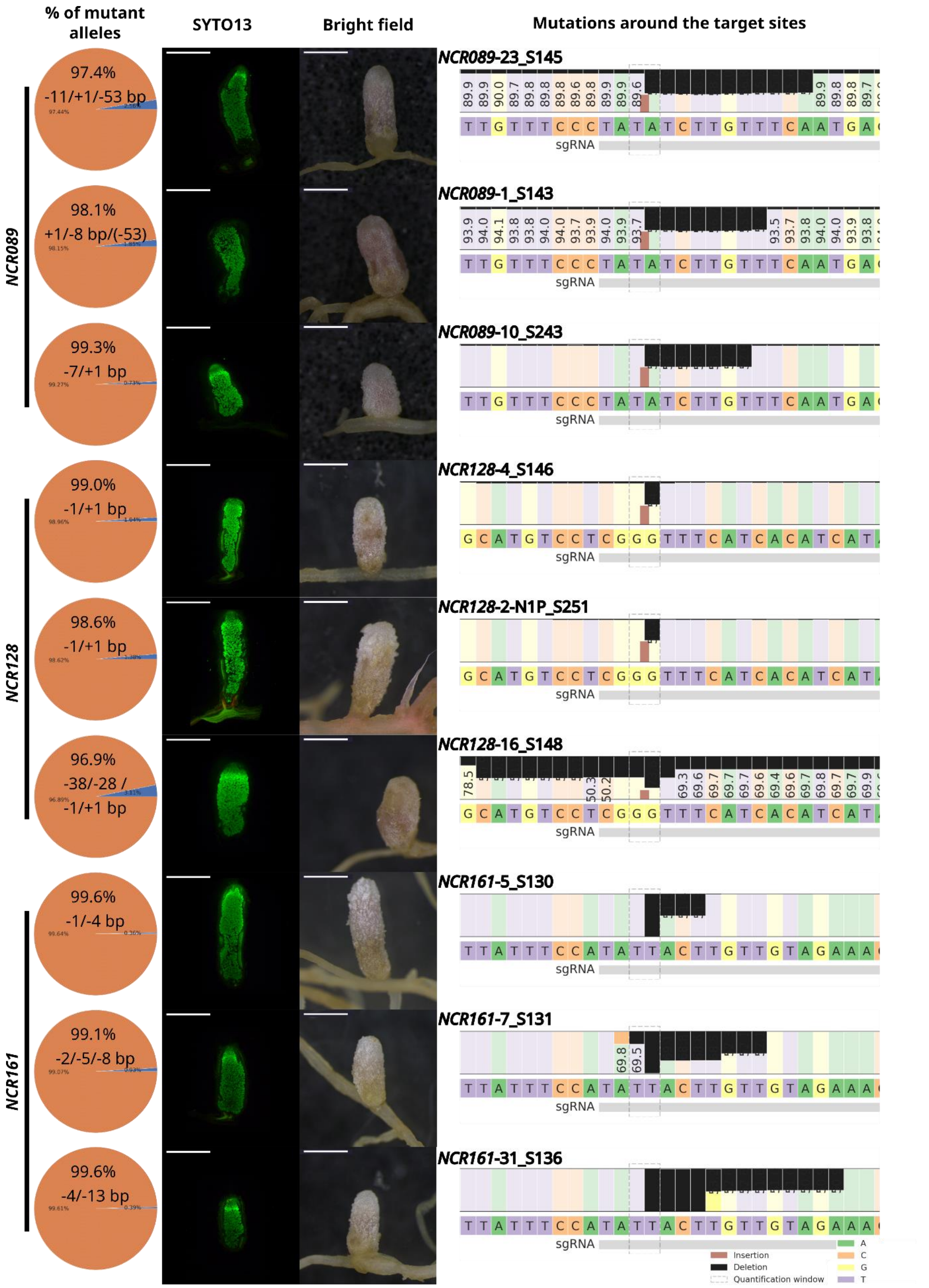


**Figure S4.** The allele sequences and the phenotype of randomly selected mutant nodules edited in genes *NCR089*, *NCR0128* and *NCR161* using the CRISPR/Cas9 method with *A. rhizogenes*-mediated hairy root transformation. Nodules were harvested 3-5 wpi with *S. medicae* WSM419 and the amplicons of targeted genes were genotyped using Illumina next generation sequencing. The sequencing result was analysed using the CRISPresso2 tool. The pie charts show the assignment of wild-type and mutant alleles of the targeted genes. All presented nodules carried frame shift mutations caused by insertions or deletions and the nature and the actual size of the mutations are listed to the right of each nodule. The nodulation phenotype of gene edited mutant roots indicates that genes *NCR089*, *NCR128* or *NCR161* are not essential for bacteroid differentiation or persistence in the symbiotic interaction between *M. truncatula* 2HA and *S. medicae* WSM419. Black and brown rectangles show deleted and inserted nucleotides, respectively.

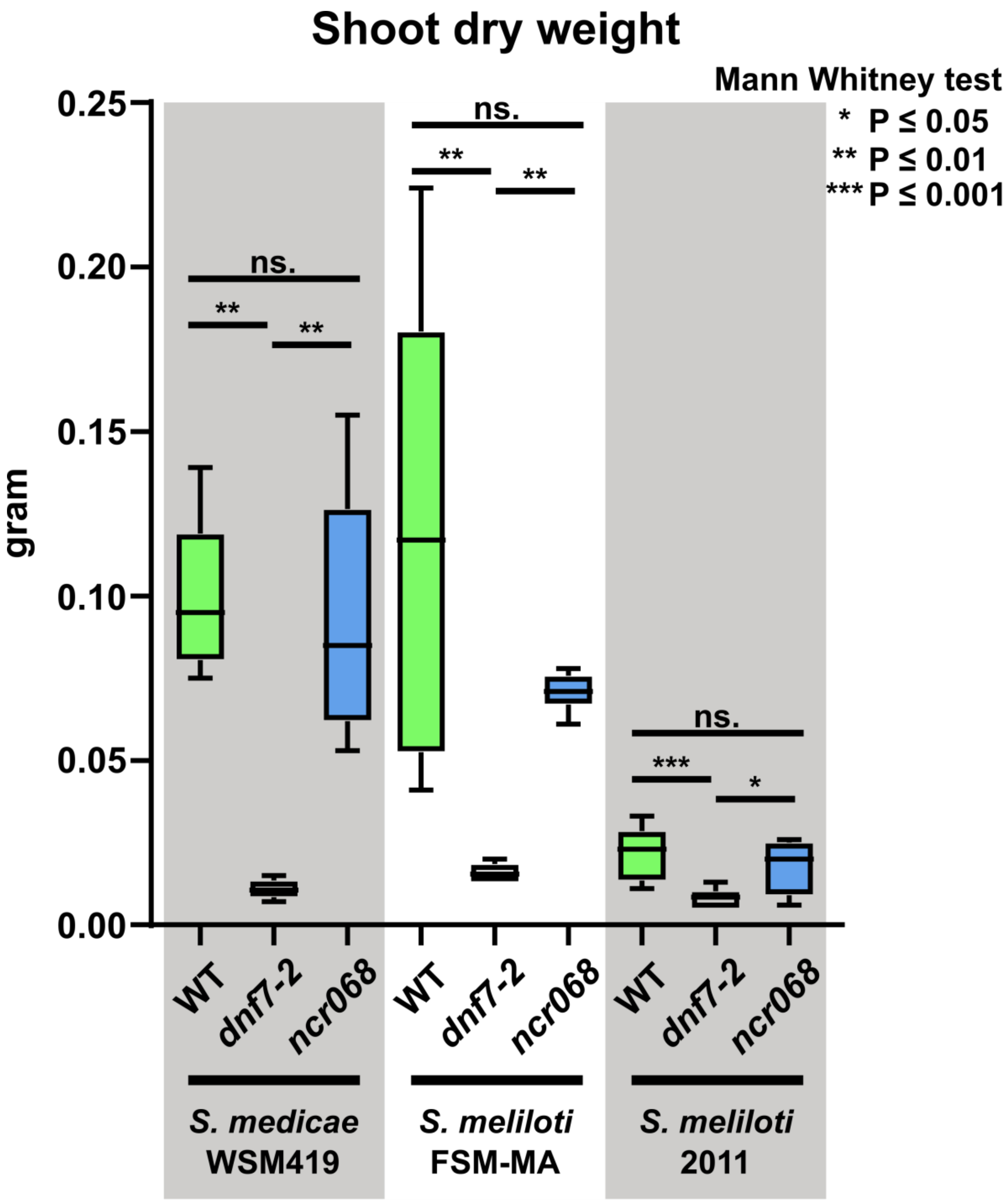


**Figure S5:** Shoot dry weight of *ncr068* mutant (T1) plants compared to wild-type (WT) and NCR169 deficient (*dnf7-2*) plants 5 wpi with three different rhizobia strains. The number of asterisks denotes significant differences at different levels (unpaired T-test). Error bars represent SE: ns.- no significant difference

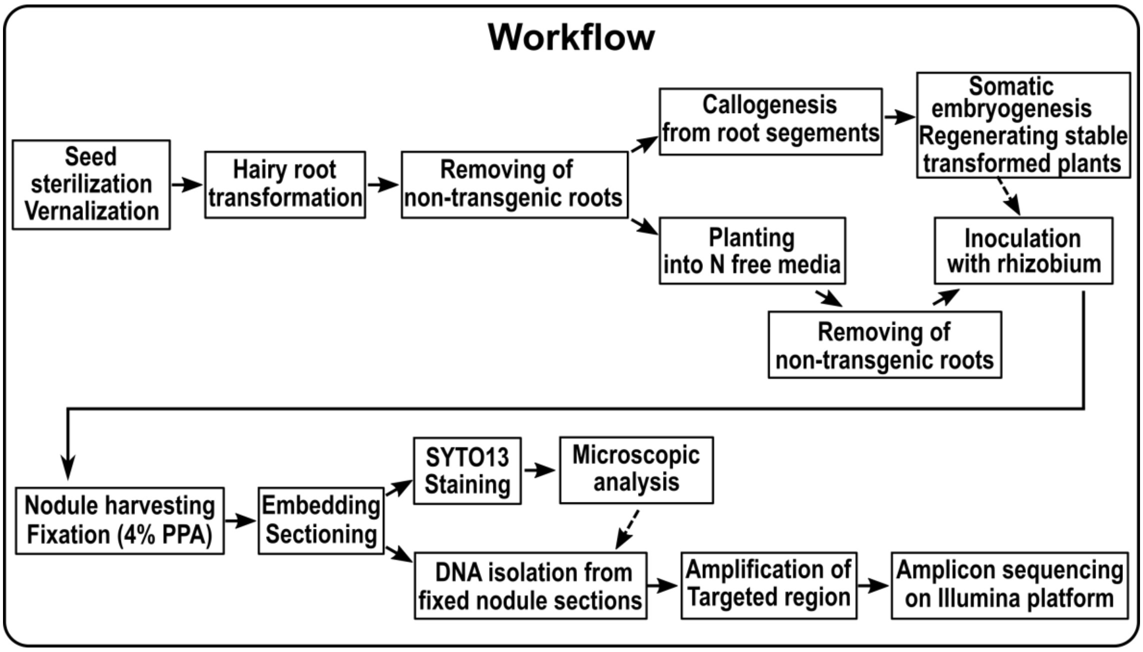


**Figure S6.** Schematic workflow of generation and analysis of induced mutations in selected *NCR* genes using the CRISPR method and *A. rhizogenes*-mediated hairy root transformation.
